# Predation risk and resource abundance mediate foraging behaviour and intraspecific resource partitioning among consumers in dominance hierarchies

**DOI:** 10.1101/364182

**Authors:** Sean M. Naman, Rui Ueda, Takuya Sato

## Abstract

Dominance hierarchies and unequal resource partitioning among individuals are key mechanisms of population regulation. The strength of dominance hierarchies can be influenced by size dependent trade-offs between foraging and predator avoidance whereby competitively inferior subdominants can access a larger proportion of limiting resources by accepting higher predation risk. Foraging-predation risk trade-offs also depend on resource abundance. Yet, few studies have manipulated predation risk and resource abundance simultaneously; consequently, their joint effect on resource partitioning within dominance hierarchies are not well understood. We addressed this gap by measuring behavioural responses of masu salmon to experimental manipulations of predation risk and resource abundance in a natural temperate forest stream. Responses to predation risk depended on body size such that larger dominants exhibited more risk-averse behaviour (e.g., lower foraging and appearance rates) relative to smaller subdominants after exposure to a simulated predator. The magnitude of this effect was lower when resources were elevated, indicating that dominant fish accepted a higher predation risk to forage on abundant resources. However, the influence of resource abundance did not extend to the population level, where predation risk altered the distribution of foraging attempts (a proxy for energy intake) from being skewed towards large individuals to being skewed towards small individuals after predator exposure. Our results imply that size dependent foraging-predation risk trade-offs can mediate the strength of dominance hierarchies by allowing competitively inferior subdominants to access resources that would otherwise be monopolized.

**Author Contributions:** SN, TS, and RU designed the study and performed the fieldwork; SN analyzed the data and wrote the manuscript with input from all authors.

## INTRODUCTION

Social dominance hierarchies and the maintenance of unequal resource partitioning among individuals are key mechanisms of population regulation and stability (Hassell 1978, Lomnicki 1988). The strength and stability of dominance hierarchies depends on the behavioural mechanisms mediating intraspecific competition (Weir and Grant 2004). Predators and resource abundance appear to have particularly important roles in this context given that many animals face a trade-off between maximizing resource intake while minimizing mortality risk (Werner and Gilliam 1984), and that the optimum of this trade-off can vary among individuals as a function of body size and social status (Lima and Dill 1990).

Size or status dependent foraging-predation risk trade-offs are often attributed to the asset protection principal (Clark 1994), which posits that larger individuals should be more risk-averse than their smaller conspecifics due to their higher accumulated fitness ‘assets’ and the diminishing energetic return for a given foraging intake with increasing body size. For consumers in dominance hierarchies, this implies that reduced foraging rates by larger dominants in the presence of predators could allow smaller subdominants to access resources that would otherwise be monopolized (Reinhardt 1999, Catano et al. 2016). As a result, predation risk should shift the distribution of resources from being highly skewed towards a small number of dominant individuals to being more evenly distributed.

The shape of foraging-predation risk trade-offs is also influenced by variation in resource abundance, which could further mediate intraspecific competition and subsequent resource partitioning among individuals (Gruber et al. 2016). Theory predicts that with increasing resources, consumers should decrease their foraging activity under predation risk (i.e., be more vigilant) due to the lower marginal benefit of food intake, which should be further reduced with increasing body size (Brown 1988, Olsson et al. 2002). Consequently, elevated resources should exacerbate the effects of predation risk on dominance hierarchies, further widening foraging opportunities for subdominants and reducing resource monopolization. However, an alternative prediction emerges if consumers are conditioned to feast or famine conditions associated with pulsed resources (Armstrong and Schindler 2011). In this case, the marginal value of foraging may be higher when resources are abundant (Higginson et al. 2012), leading to higher foraging rates and agonistic interactions. Consequently, elevated resources would reduce foraging opportunities for subdominants and dampen or even reverse the influence of predation risk on resource monopolization.

Despite broad support for foraging-predation risk trade-offs as drivers of intraspecific competition, our ability to predict the specific outcomes of these factors at the population level is currently limited as surprisingly few studies have manipulated both predation risk and resource abundance simultaneously (for exceptions see: Kotler, Brown, & Bouskila, 2004; Matassa & Trussell, 2014; Morosinotto, Villers, Varjonen, & Korpimäki, 2017). Further, these cases are often highly controlled experiments, where resource abundance levels and foraging motivation are tightly regulated. While this is preferable for teasing apart the specific mechanisms underlying behaviour, results may not necessarily translate to resource partitioning in natural populations, where foraging opportunities and among-individual feeding motivation can be highly variable (Yang et al. 2008, 2010). Moreover, inferences into social interactions may be complicated by behavioural artefacts introduced in more controlled settings (Sloman and Armstrong 2002). Therefore, conducting behavioural studies *in situ* is key to thoroughly understand the role that foraging-predation risk trade-offs actually play in natural systems.

Here we present the results of a field experiment testing how consumers (red-spotted masu salmon: *Oncorhynchus masou ishikawae*) respond to manipulations of predation risk and resource abundance in a temperate forest stream. Stream salmonids are an ideal taxon to test these ideas given their behaviour is easily observable in the field (Nakano 1995), they often exhibit strong dominance hierarchies where larger dominant individuals exclude smaller subdominants from the most profitable foraging territories (Nielsen 1992, Weir and Grant 2004), and they frequently experience significant predation risk from terrestrial predators (Hoeinghaus and Pelicice 2010, Harvey and Nakamoto 2013)

In this study, we first tested whether behavioural responses by masu salmon to predation risk varied with body size and whether this response was mediated by resource abundance. Based on the asset protection principle (Clark 1994), we predicted that riskier behaviour, defined as higher foraging rates after predator exposure, would decline with body size. We further predicted that these effects would be magnified when resources were elevated due to a lower marginal benefit of energy intake for dominant individuals. Although absolute body size may be the ultimate driver of foraging-predation risk trade-offs, behavioural responses may be strongly mediated by social status (dominant vs. subdominant) among directly interacting individuals (Gotceitas and Godin 1991). Thus, we further tested how behavioral responses to predation risk and resource abundance were mediated by social status within a dominance hierarchy, predicting that subdominants should disproportionately benefit from elevated resources under predation risk. The corollary of these individual-level predictions is that predation risk combined with elevated resource abundance should result in reduced resource monopolization, i.e., a more even distribution of resources. Thus, at the population level, we tested whether predation risk and resources modified the distribution of foraging attempts (a surrogate for relative energy intake) among individuals, predicting that subdominants should gain an increasingly greater share of energy with elevated resources under predation risk.

## METHODS

### Study system and logistics

Our study was conducted in Hachiman-dani stream, an upper tributary of the Arida River in Kyoto University’s Wakayama Forest Research Station in Japan. The study system is a cobble-bottomed stream draining a catchment dominated by planted conifer trees (*Cryptomeria japonica*) and native deciduous vegetation. Masu salmon and low densities of minnows (*Rhynchocypris oxycephalus jouyi*) are the only fish species present in the study area.

We conducted our study in conjunction with a large-scale field experiment testing the influence of pulsed resources on the life history of masu salmon (T. Sato unpublished). That experiment involved separating the study stream into six experimental reaches, consisting of three replicates of two resource treatments (control vs. elevated). Reaches were separated by check dams, which constrained movements of fish (< 1% individuals moved across reaches during the two-year experiment; Sato et al. unpublished data). To manipulate resource abundance, live mealworms (*Tenobrio molitor*) were added to the stream at the rate of 100 mg m^-2^ day^-1^, which corresponds to peak input rates of natural terrestrial invertebrates in temperate forest streams (e.g., Baxter, Fausch, & Saunders, 2005; Nakano & Murakami, 2001). Mealworms were dispensed by automatic fish feeders (W × H × D = 6.8 × 14.9 × 8.7 cm, ~100 mL capacity for food; EHEIM Co. Ltd.) set on wooden stakes ~1.5 m above the stream water surface placed at 10 m intervals throughout the reach. The feeders were deployed in similar habitats (i.e., a riffle upstream of a pool) so that fish had similar access to mealworms. We set each feeder to dispense mealworms four times during daylight (6:00-18:00), which mimicked the slow and haphazard manner by which salmonids encounter natural prey falling from riparian vegetation.

We quantified natural variation in ambient prey abundance by sampling aquatic and terrestrial invertebrate drift with 250 μm nets staked to the substrate (*n* = 5 per treatment reach). On two days during the experiment, three 30 minute samples were taken during daylight and pooled for a single estimate at each location. Invertebrates were identified to order, dried, and weighed in the laboratory. Drift concentrations (mg volume filtered^-1^) was converted to a total flux through each pool following Downes & Lancaster (2010) and biomass was converted to energy (kilojoules) using taxa-specific conversions (Sato et al. 2011). After accounting for drift, ambient terrestrial invertebrate input rates (~20 mg-dry mass m^-2^ day^-1^; Sato *et al*. unpublished data), and for the added mealworms, we determined that fish in elevated resource reaches had access to 7.5 times more energy on average than those in controls (Figure 1).

**Figure 1.**
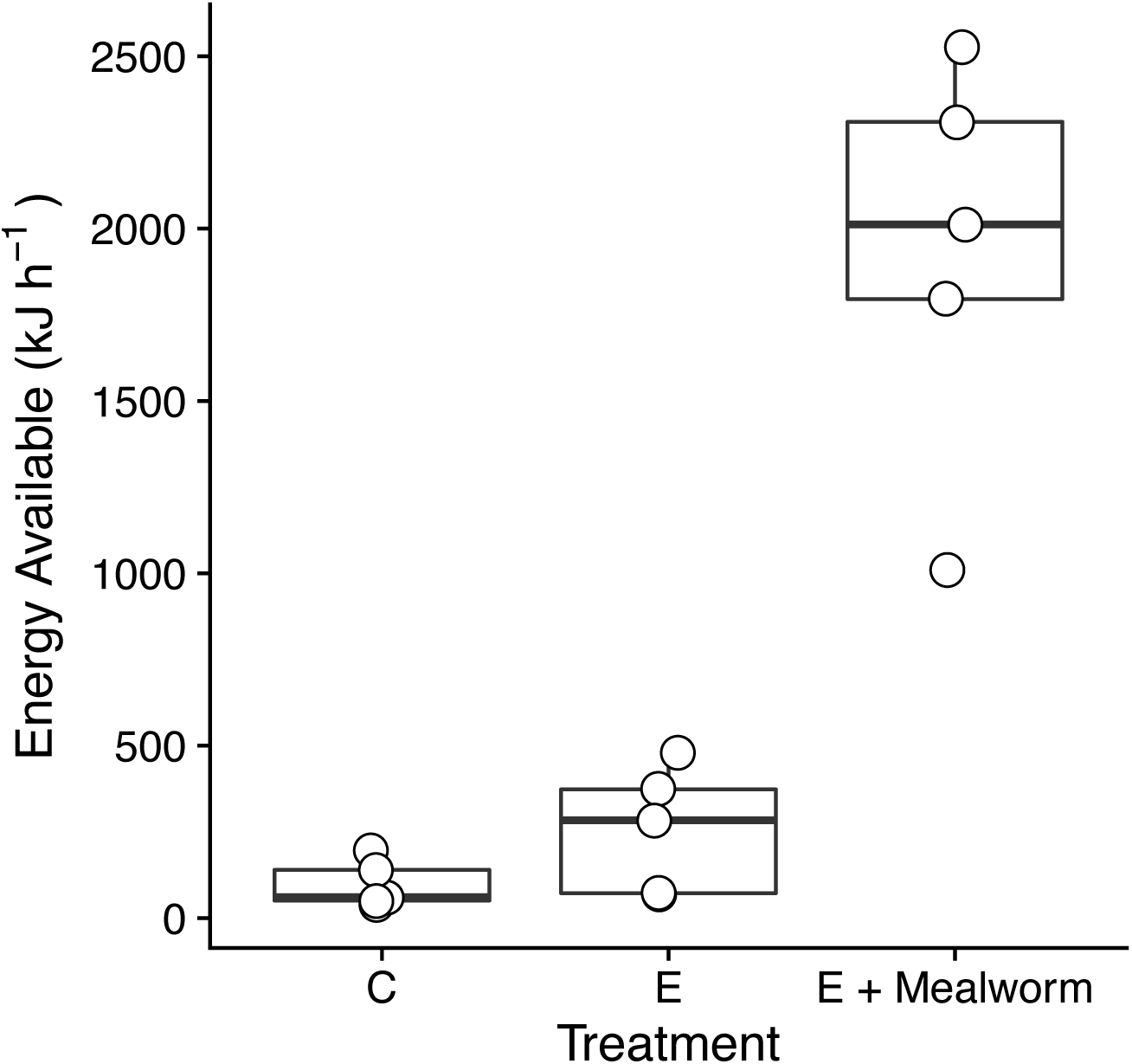
Boxplots showing the total invertebrate energy available to fish (kJ hour^-1^) in control treatments (C), elevated resource treatments before mealworm additions (E), and elevated resource treatments after mealworm additions (E + Mealworm).

We took advantage of the infrastructure from the larger experiment to examine behavioural responses to predation risk under ambient (controls) and experimentally elevated resources. Our study began one week after mealworm treatments were initiated, such that fish had sufficient time to adjust to the new prey source. Sampling of stomach contents further confirmed that fish consumed mealworms in elevated treatments (Supplement Fig S1). We first selected pools (*n* = 10 per treatment) with similar abiotic attributes and similar densities of fish (~0.5 m^-2^). Masu salmon generally forage in pool habitats and maintain strong size-based dominance hierarchies, where larger dominant fish occupy upstream positions that are more energetically profitable to intercept aquatic and terrestrial invertebrates (Fausch 1984, Nakano 1995). Social status of fish in the experimental pools was easily identified by body size and corroborated by numerous observations of aggressive interactions where dominant individuals would chase subdominants out of foraging territories (Sato and Watanabe 2014).

Observations of fish behaviour were made using underwater videography. We attached underwater video cameras (Seesii 30M 7” LCD, EYOYO Co. Ltd.) to rebar stakes anchored into each pool at locations permitting the widest possible field of view. Two cameras were used for wide or irregularly shaped pools to allow all fish to be visible. Cameras were left overnight after installation and then were connected to viewing monitors, which were positioned out at least 15 m away from the pool where fish behaviour could be observed while minimizing any artefacts of human presence.

Predator simulations consisted of a decoy of a crow, which was fashioned to a fishing line that we tied across each pool 3 weeks prior. Lines were tied at a sufficient distance from the pool such that the investigator would not disturb fish when releasing them. There were no noticeable effects of moving the overhead line on fish behaviour. Before initiating predator simulations, we ensured fish were present in the pools then began recording video for at least 30 minutes. Then, one investigator stealthily attached the bird decoy to the line and released it while a second observed fish behaviour through the monitor to ensure no artefacts were present. The bird was rigged to fly over the pool from upstream to downstream at roughly 15-degree angle at a speed of 0.5 body lengths sec^-1^, making brief contact with the water. We marked the time of the exact moment that fish reacted to the predator simulation, then proceeded to record footage for at least one hour. Technical difficulties with cameras resulted in recording times differing slightly, thus subsequent calculations are standardized by recording time. All predator simulations occurred between 10:00 and 14:00 to minimize effects of diel variation in fish activity.

The predator simulation was designed to replicate the behaviour of several avian predators in the study area, including two kingfishers (*Halcyon coromanda* and *Megaceryle lugubris*) and brown dippers (*Cinclus pallasii*). Several opportunistic observations suggested fish responses to the decoy were qualitatively similar to their responses to these predators (S. Naman and T. Sato personal observation). While there is some evidence for size-biased avian predation on salmonids (Miyamoto et al. 2018), previous observations of visible injury to captured fish suggest that predation risk is not strongly size biased within the range of fish sizes in our experiment (55-162 mm; T. Sato unpublished data).

### Behaviour observations

We quantified changes in foraging behaviour for each observed fish in response to predation risk and resource abundance several different ways following Nakano (1995). First, we defined the appearance rate (AR) as the proportion of time a fish was visible in the pool relative to the total footage recorded. Second, we define the frequency of foraging attempts (FFA) as the number of foraging attempts per minute that a given fish was visible. Third, we define the actual foraging rate (AFR) as the product of AR and FFA. We also quantified the number of aggressive interactions across all fish in a given pool during each observation period. While we did not explicitly quantify their direction of initiation or outcome, nearly all interactions involved dominants chasing or charging subdominants.

We visually estimated fish size using ceramic tiles with known dimensions placed in the pool during camera installation. This technique was validated by capturing fish by backpack electrofishing in a subset of pools immediately after observations, which indicated we were able to correctly estimate size within 5 mm on average (*n* = 9, *r* = 0.70). Individual fish within each pool were generally identifiable by their relative size and an ongoing mark-recapture experiment has demonstrated an extremely high site fidelity of fish to individual pools in the study site (Sato and Watanabe 2014). Thus, we were confident that fish returning to a given pool after predator exposure were the same fish that were originally there. In several cases there was uncertainty in this regard and these observations were not incorporated in subsequent analysis. Altogether, behavioural observations of 34 fish were analyzed.

### Statistical Analysis

We used generalized linear mixed effects models (GLMM) to test whether body size influenced behavioural responses to predation risk and resource abundance. For these analyses, the response variable was the change in a given foraging metric (AR, FFA, AFR) from before to after exposure to the predator decoy. The predictors were body size, resource abundance (ambient control or elevated), and their interaction. Each pool was treated as a random factor. We fit models using the R package *glmer* (Bates et al. 2015) with a Gaussian error distribution and tested the significance of each predictor using sequential likelihood ratio tests. For significant predictors (*P* < 0.05) we computed 95% confidence intervals using a parametric bootstrap (*n* = 10,000 iterations).

We tested the difference in foraging rates between subdominants and dominants in each pool using a similar GLMM approach, but in this case resource abundance, time (before-after predator exposure) and their interaction were predictors. Fish ID was used as a random factor to account for repeated measures on the same fish. We initially included pool as another random factor (where fish ID is nested within pool) but this did not improve model fits and resulted in poor parameter convergence so only fish ID was retained; moreover, only two pools had more than two fish. We tested for main treatment effects as previously described and also examined whether 95% confidence intervals in each group (e.g., control-before, elevated-after etc.) overlapped zero, which would indicate that foraging rates of subdominants did not differ from dominants on average. We tested treatment effects on the frequency of aggressive interactions using a GLMM with a quasi-Poisson distribution. All data were analyzed in R v. 3.2.0 (R Core Development Team). Model diagnostics for *glmer* were evaluated with residual simulations using the package DhArma (Hartig 2016).

To determine the effects of predation risk and resource abundance on energy distribution among individuals and the extent of resource monopolization, we compared the frequency distributions of the total foraging attempts observed across body lengths (mm) before versus after predator exposure and among resource treatment combinations. While foraging attempts as a metric of absolute energy intake is inappropriate given many attempts are likely unsuccessful (Neuswanger et al. 2014), we assume it is a reasonable proxy for the *relative* energy intake among individuals. We specifically tested the prediction that predation risk and resource abundance should lead to more positively skewed foraging attempt-body size distributions, indicating a higher relative energy intake by smaller individuals. Skewness was determined by fitting Gamma probability functions to scaled frequency distributions by maximum likelihood using the R package *fitdistplus* (Delignette-Muller and Dutang 2015), deriving estimates and 95% bootstrapped confidence intervals of shape parameters (α), then computing skewness using the formula: 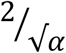

We inferred if skewness differed between two distributions if 95% confidence intervals did not overlap. Higher skewness values indicate a longer right tail; thus, an increase in skewness following predator exposure would indicate a shift in energy distribution to smaller individuals.

## RESULTS

Fish reacted strongly to predator simulations, generally by burst swimming out of the pool or moving close to the substrate. As predicted, individual-level foraging rates following predator exposure were inversely related to body size (Figure 2), with all foraging metrics declining for the largest fish while increasing for smaller fish. However, this relationship was not mediated by resource abundance as the interaction between resource abundance and fish size was not significant for any of the foraging metrics we examined (Supporting Information, Table S1).

**Figure 2.**
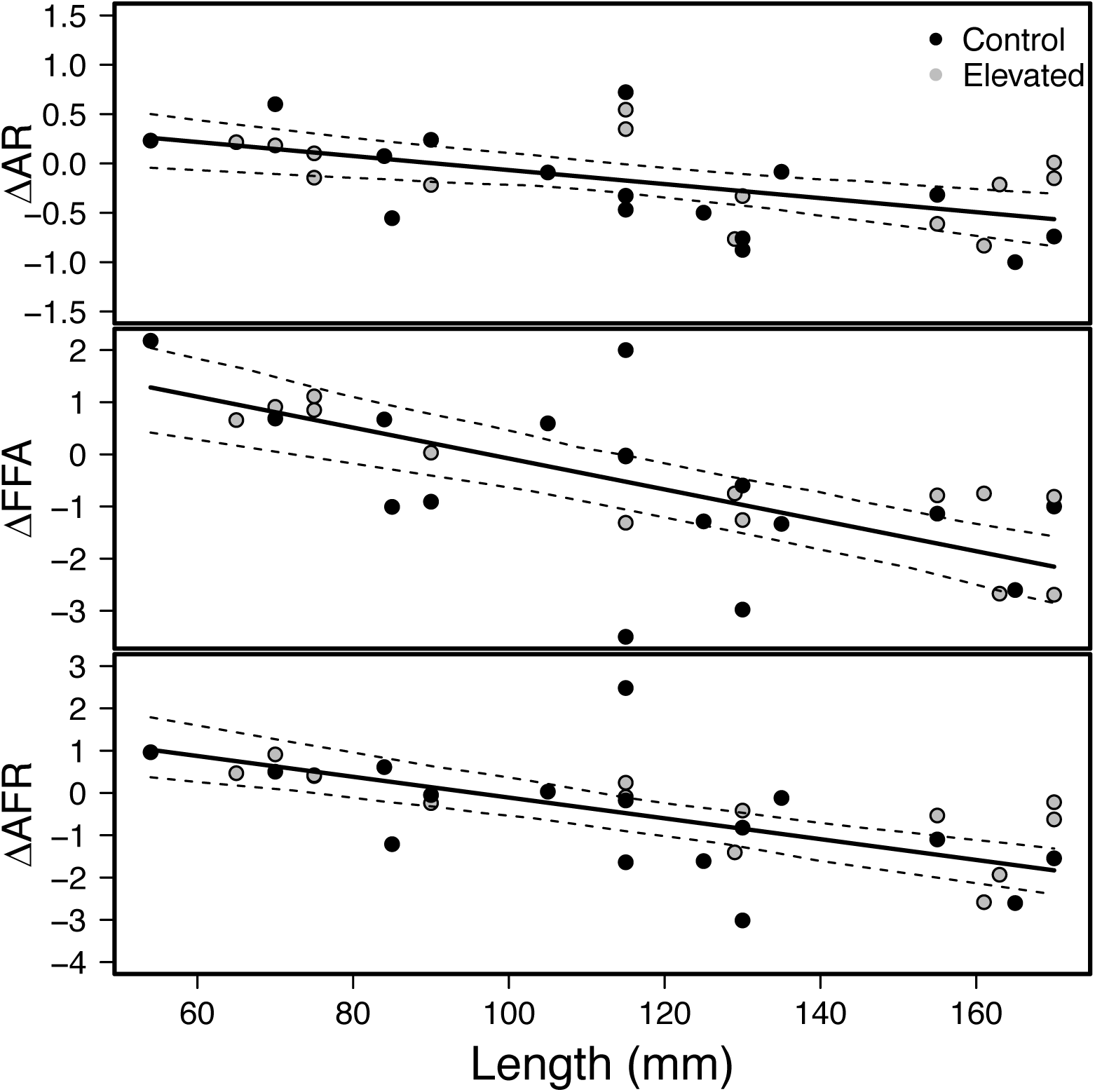
The change in appearance rate (AR), frequency of foraging attempts (FFA), and actual foraging rate (AFR) for fish in elevated and control resource treatments from before to after predator exposure. Solid lines indicate predictions from GLMM models and dashed lines indicate 95% confidence intervals. Note the differing scales of y-axes.

When foraging behaviour was examined in the context of dominant-subdominant interactions within individual pools, effects of both predation risk and resources were evident, albeit contrary to our predictions. Before predator exposure, appearance rates of subdominants were equal to dominants in pools with elevated resources but lower than dominants in control pools with ambient resources (Figure 3). This pattern changed after predator exposure such that appearance rates were still equal with elevated resources but subdominants exceeded dominants in controls, i.e., there was a significant resource-time interaction (Table S1). Foraging attempt frequency was lower for subdominants in both resource treatments before predator exposure but changed after exposure such that subdominants foraged more frequently than dominants in controls but not in elevated resource treatments where foraging frequency was equivalent (Figure 3). Actual foraging rate (i.e., appearance rate × foraging attempt) followed a similar pattern as the frequency of foraging attempts, which had larger effect sizes than appearance rate (Table 1). These changes in foraging metrics appeared to be primarily driven by changes in the absolute foraging rates of dominant fish, whereas subdominant foraging rates were relatively constant (Figure 3)

**Figure 3.**
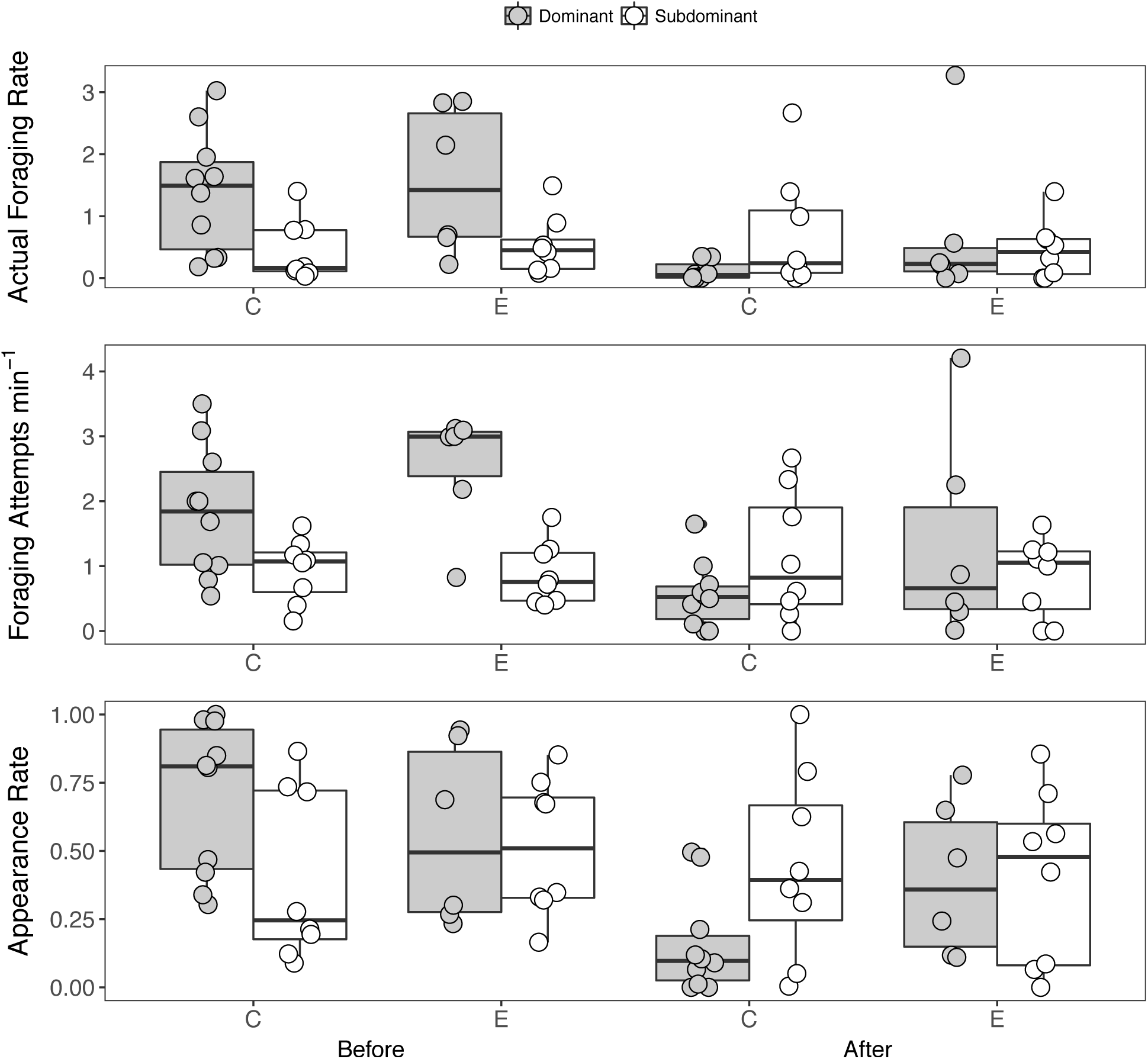
Foraging metrics of subdominants and dominants in control (C) and elevated resource (E) treatments, before and after predator exposure. Boxes are 25% and 75% quantiles and overlaid points are raw values. Note that y-axis values are on different scales.

**Table 1.** GLMM parameter estimates and 95% confidence intervals for the difference between subdominant and dominant foraging rates. Cases where confidence intervals did not overlap zero are highlighted in bold.

| Treatment | Estimate | 5% CI | 95% CI |
| --- | --- | --- | --- |
| *Appearance Rate* |  |  |  |
| Control-Before | **-0.23** | **-0.46** | **-0.02** |
| Elevated-Before | 0.03 | -0.20 | 0.26 |
| Control-After | **0.37** | **0.15** | **0.60** |
| Elevated-After | 0.13 | -0.09 | 0.36 |
| *Foraging Frequency* |  |  |  |
| Control-Before | **-0.78** | **-1.40** | **-0.17** |
| Elevated-Before | **-1.76** | **-2.38** | **-1.13** |
| Control-After | 0.61 | 0.00 | 1.23 |
| Elevated-After | -0.01 | -0.63 | 0.60 |
| *Actual Foraging Rate* |  |  |  |
| Control-Before | **-0.76** | **-1.34** | **-0.18** |
| Elevated-Before | **-0.83** | **-1.40** | **-0.25** |
| Control-After | **0.67** | **0.08** | **1.24** |
| Elevated-After | 0.26 | -0.31 | 0.83 |

The frequency of aggressive interactions among masu salmon in each pool responded to both predation risk and resource abundance, but there was no evidence for an interaction [Time (before/after-predation risk): *χ*^2^ = 19.2, *P* < 0.001; Resources: *χ*^2^ = 5.3, *P* = 0.02; interaction: P = 0.7]. Aggressive interactions decreased by 9-fold on average from before to after predator exposure and were 15 and 6-fold higher in elevated resource treatments relative to controls before and after predator exposure respectively.

Examining the frequency distribution of total foraging attempts across body size largely supported our prediction that predation risk should redistribute resources to smaller individuals (Figure 4). Specifically, skewness was higher after relative to before predator exposure when resource treatments were pooled together, indicating a shift toward smaller individuals [before: 0.53 (95% CIs, 0.50-0.57); after: 0.63 (95% CIs 0.58-0.69)]. Distributions appeared to differ when resource treatments were separated, with foraging attempts being more skewed towards larger individuals before predator exposure, and more evenly distributed after predator exposure when resources were elevated (Figure 4). However, confidence intervals around skewness estimates overlapped between all resource-time treatment combinations.

**Figure 4.**
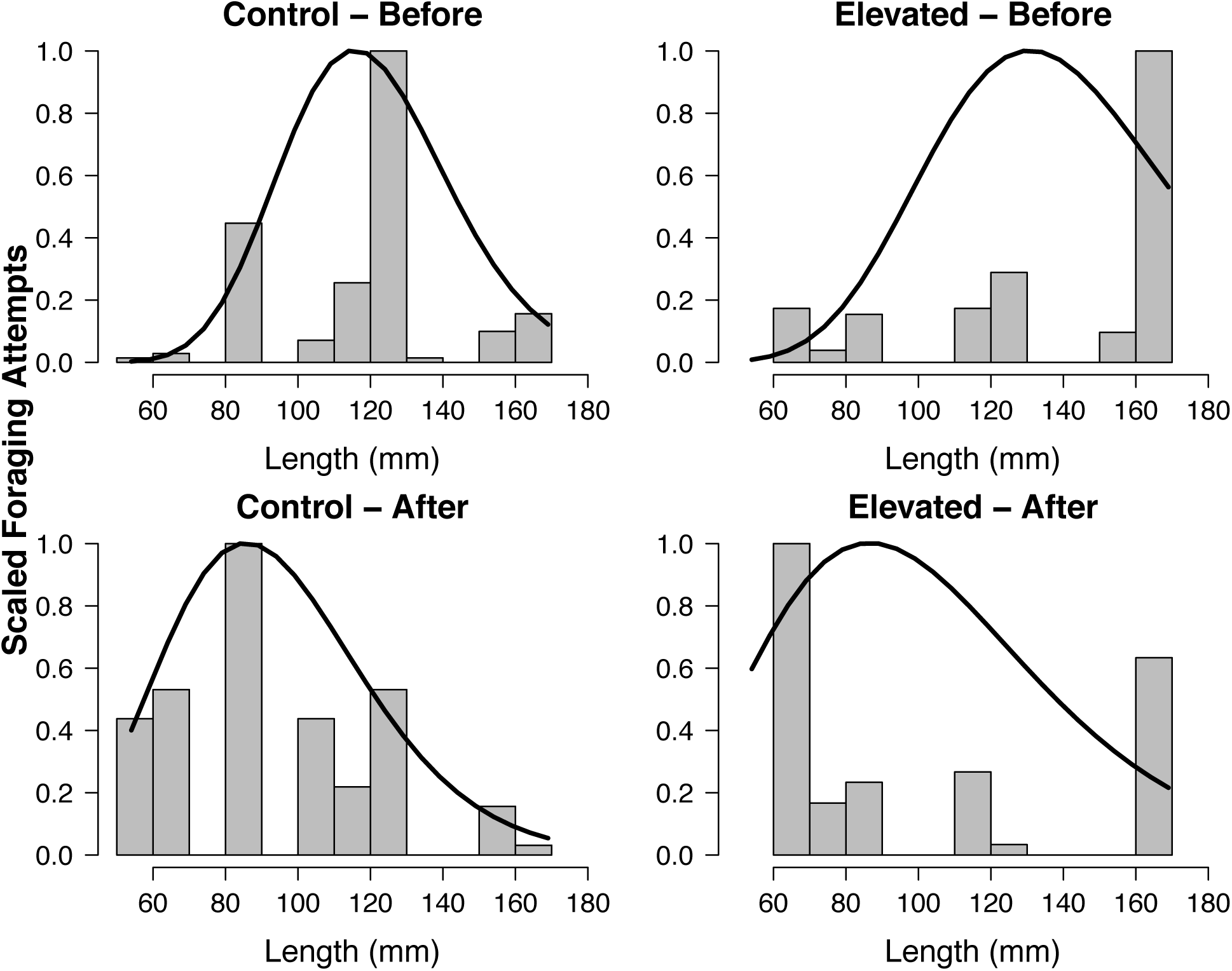
The distribution of relative foraging attempts among different body sizes in each control and elevated resource treatments, before and after predator exposure. Distributions are scaled to a maximum of 1. Black lines represent gamma probability density functions fit to the data in each group.

## DISCUSSION

Dominance hierarchies and unequal partitioning of resources are well known to be key mechanisms of population regulation (Lomnicki 1980, Keeley 2001) and size-dependent foraging-predation risk trade-offs have been observed over a wide range of taxa including salmonids (Reinhardt 2002, Kotler et al. 2004, Morosinotto et al. 2017). Our experiment is relatively unique in its integration of these concepts insofar as it incorporated simultaneous manipulations of predation risk and resource abundance, and explicitly tests the consequences of individual-level behaviour on intraspecific resource partitioning at the population level.

Together, our results suggest that body size and social status drive responses to predation risk and resource abundance among individuals, which in turn influences the extent of resource monopolization, and ultimately the distribution of energy within populations. While we are limited in our ability to disentangle the specific mechanisms underlying these findings, the patterns we observed are arguably strengthened when viewed against the background variation associated with wild populations in natural conditions.

Consistent with the asset protection principle, behavioural responses to predation risk varied with absolute body size, with larger individuals more risk averse relative to their smaller conspecifics. The effects of resource abundance on these responses were more nuanced and appeared to be mediated by social status among fish within each pool; they were also contrary to predictions. In particular, dominant fish decreased their foraging rates in both treatments following predator exposure but less so when resources were elevated. This contradicts our expectation that consumers should be more vigilant due to a lower marginal benefit of foraging on abundant resources (McNamara and Houston 1986, Kotler et al. 2004). Instead, this result may support the alternative hypothesis, where feast-or-famine conditions lead consumers to accept higher predation risk when resources are abundant in order to meet their energetic demands (Higginson et al. 2012). In our system, masu salmon experience long periods of resource limitation due to large-scale deforestation followed by a monotonic conifer plantation, leading to depressed growth (Sato unpublished data). Thus, we speculate that mealworm additions may have represented a crucial foraging opportunity where increased foraging under risky conditions was necessary to meet energetic demands over longer timescales (e.g., seasonal growing periods).

Foraging metrics for subdominant fish were largely consistent across all treatments. Consequently, the foraging rates of subdominants *relative* to dominants increased following predator exposure, especially in control treatments where dominants were more vigilant. This suggests that subdominants accepted a greater predation risk relative to dominants in controls, which may have been due to subdominants having lower energetic status such that avoiding starvation necessitated riskier behaviour (Lima and Bednekoff 1999). However, subdominant foraging rates were unchanged in elevated resource treatments suggesting they did not accept a proportionally higher level of risk with a higher energetic return. One explanation could be that subdominants were sufficiently satiated such that their energetic demands could be met without increasing their foraging rates. We cannot rule out this possibility; however, it appears unlikely in light of the strong resource limitation these fish experienced prior to mealworm additions (Sato Unpublished data). Alternatively, this inconsistency could potentially be explained by aggressive interactions, which were significantly higher when resources were elevated. Agonistic interactions are energetically costly and reduce foraging time (Puckett and Dill 1985, Metcalfe 1986, Mathot and Dingemanse 2015), thus may have mediated these responses.

Inferences into the mechanisms underlying individual-level behaviour should be tempered by several caveats inherent in our design. First, body size and social status likely covary with energetic state, which we did not explicitly account for and may be the ultimate driver of short-term behavioural responses (Gotceitas and Godin 1991). Second, we cannot completely rule out the possibility that perceived predation risk varied with body size (Miyamoto et al. 2018). While these uncertainties cannot be resolved with the data at hand, they do not change any of the conclusions of our study so much as offer alternative mechanisms for them. Ultimately, predation risk and resource abundance still altered foraging behaviour and intraspecific interactions, leading to a shift in population-level resource distribution toward smaller individuals. Thus, our study as a whole provides an important *in situ* demonstration of state-dependent behavioural responses to predation risk and resource abundance, and its potential consequences for populations.

While our inferences are constrained by the short duration of our study, the redistribution of resources we observed may further affect population dynamics over longer time scales. Theory of density-dependent population regulation predicts that more equal partitioning of resources should result in unstable population dynamics relative to skewed distributions where resources are monopolized by a small number of competitively superior individuals (Lomnicki 1980, Johst et al. 2008). The extent that population dynamics are related to short term behavioural changes that modify dominance structure is an important question, especially with regards to territorial taxa like salmonids. To further bridge the gaps between short-term behaviour and long-term population processes, the following issues should be addressed: (1) the frequency that consumers experience predation risk over short and long time scales in natural systems; (2) whether consumers can adaptively respond to temporal variability in predation risk and resource abundance via memory and behavioral adjustment (Lima and Dill 1990, Armstrong and Bond 2013); and (3) whether these adaptive responses (if any) shape the foraging-predation risk trade-offs and consequent resource partitioning.

## ACKNOWLEDGMENTS

We thank the staff at the Wakayama Forest Research Station for providing accommodation, logistical support, as well as keen insights into experimental design. Shin Ukaji, and Ryosuke Tanaka, and Torio were invaluable field assistants. A UBC Go Global fellowship supported travel for SN. TS was supported by JSPS KAKENHI Grant-in-Aid for Scientific Research (Grant Number: 15H04422).

## PERMITS

The experiment was conducted in accordance with the regulations for animal experiments at Kobe University. Electrofishing was conducted according to regulations of Inland Fisheries Adjustment of Wakayama Prefecture with the permission of the Department of Agriculture, Forestry and Fisheries, Wakayama Prefecture.

## SUPPORTING INFORMATION

Figure S1: Stomach contents of masu salmon in each resource abundance treatment.

Table S1: Summary statistics for GLMM analysis of the effects of body size and social status on foraging rates.

## Supporting Information

**Table S1:** GLMM results for the change in foraging rates from before to after predator exposure and the difference in foraging rates between subdominant and dominant individuals. *χ*^2^ and P values are from sequential likelihood ratio tests. R^2^_Marginal_ indicates the proportion of variation explained by only fixed effects; R^2^_conditional_ indicates the variance explained by fixed and random effects.

| Model Term | $\chi^2$ | P | $R^2_{\text{Marginal}}$ | $R^2_{\text{Conditional}}$ |
| --- | --- | --- | --- | --- |
| <u>Change in Foraging Rates After Predator Exposure</u> |  |  |  |  |
| *Actual Foraging Rate* |  |  | 0.37 | 0.62 |
| Body Size | 15.69 | 0 |  |  |
| Resources | 0.41 | 0.52 |  |  |
| Body Size x Resources | 0.14 | 0.7 |  |  |
| *Foraging Frequency* |  |  | 0.36 | 0.60 |
| Body Size | 16.85 | 0 |  |  |
| Resources | 0.11 | 0.73 |  |  |
| Body Size x Resources | 0.18 | 0.75 |  |  |
| *Appearance Rate* |  |  | 0.36 | 0.49 |
| Body Size | 11.26 | 0 |  |  |
| Resources | 1.2 | 0.28 |  |  |
| Body Size x Resources | 1.5 | 0.21 |  |  |
| <u>Subdominant-Dominant Foraging Differences</u> |  |  |  |  |
| *Actual Foraging Rate* |  |  | 0.38 | 0.39 |
| Resources | 0.00 | 0.95 |  |  |
| Time | 18.61 | 0.00 |  |  |
| Resources x Time | 3.06 | 0.08 |  |  |
| *Foraging Frequency* |  |  | 0.50 | 0.56 |
| Resources | 2.77 | 0.10 |  |  |
| Time | 23.87 | 0.00 |  |  |
| Resources x Time | 0.29 | 0.59 |  |  |
| *Appearance Rate* |  |  | 0.31 | 0.31 |
| Resources | 0.31 | 0.58 |  |  |
| Time | 10.20 | 0.00 |  |  |
| Resources x Time | 10.11 | 0.00 |  |  |

**Figure S1.**
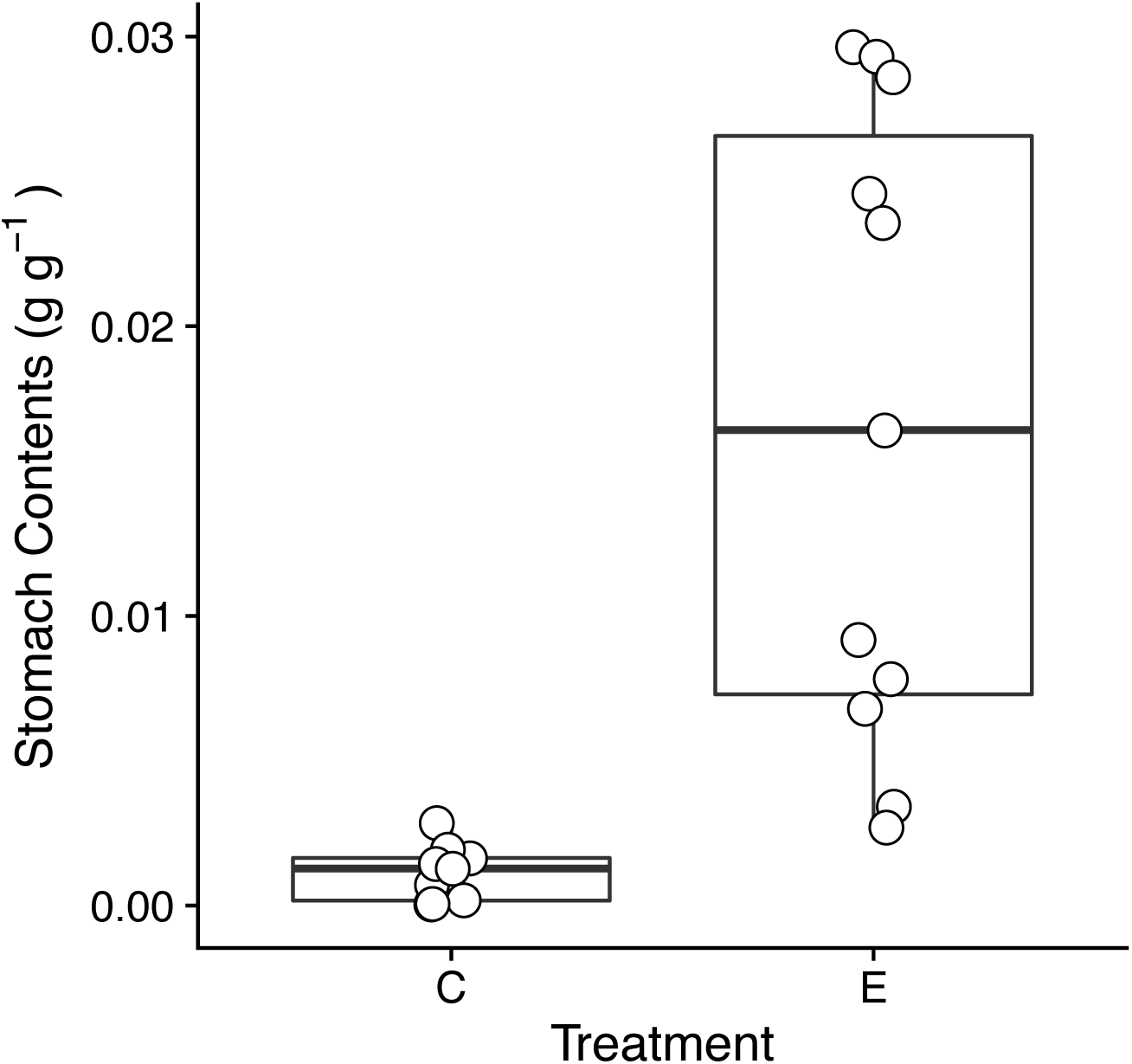
Mass specific stomach content biomass (g dry mass invertebrates · g wet mass fish^-1^) of masu salmon in control (C) and elevated resource (E) treatments. Stomach contents were collected with gastric lavage two weeks prior to the experiment. Contents were stored in 75% ethanol, identified to order, oven dried at 60° C, and weighed in the laboratory. Mealworms constituted ~70% of the stomach content biomass in elevated treatments on average.

